# Viscoelastic properties of ECM-rich embryonic microenvironments

**DOI:** 10.1101/853044

**Authors:** Zsuzsa Akos, Dona G. Isai, Sheeja Rajasingh, Edina Kosa, Saba Ghazvini, Prajna Dhar, Andras Czirok

## Abstract

The material properties of tissues and their mechanical state is an important factor during development, disease, regenerative medicine and tissue engineering. Here we describe a microrheological measurement technique utilizing aggregates of microinjected ferromagnetic nickel particles to probe the viscoelastic properties of embryonic tissues. Quail embryos were cultured in a plastic incubator chamber located at the center of two pairs of crossed electromagnets. We estimate the Young’s modulus of the ECM-rich region separating the mesoderm and endoderm in Hamburger Hamilton stage 6-10 quail embryos as 300±100 Pa. We found a pronounced viscoelastic behavior consistent with a Zener (standard generalized solid) model. The viscoelastic response is about 45% of the total response, with a characteristic relaxation time of 1.3 sec.

## 2 introduction

Tissues are physical bodies, thus their formation necessarily involves controlled generation and relaxation of mechanical stresses (Preziosi et al., 2010). Tissue cells are known to generate mechanical stresses by actin-myosin contractility, specifically relying on non-muscle Myosin II, with upstream regulators coordinated through a spatial and temporal activity of rho GTPases such as RhoA (Ridley et al., 2003). The relaxation of mechanical stresses involves the disruption of cell-cell connections, often accompanied by changes in cell neighbors (Forgacs et al., 1998; Smutny et al., 2017; Petridou et al., 2019). While this process is less understood on the molecular level than acto-myosin contractility, the spatio-temporal regulation for both force generation and relaxation are equally important to shape the embryonic tissues. Embryonic tissues are thus plastic, with their stress-free shapes deforming through the development process.

A cell-resolved mechanism underlying tissue plasticity was first resolved in flies, where studies indicated a pulsatile, ratchet-like contraction mechanism (Martin et al., 2009). Thus, instead of a uniformly distributed contractile activity across the tissue, individual cells were observed to undergo (asynchronously) a repeating cycle of contraction, stiffening and relaxation by cytoskeletal rearrangements. The pulsatile nature of tissue movements is also evident in the ECM displacements recorded within avian embryos (Szab et al., 2011).

While measures for tissue deformation (strain) became recently possible to obtain during development (Rozbicki et al., 2015), estimates for tissue stress and material properties are still very challenging to determine. A FRET-based molecular sensor has been recently developed (Meng and Sachs, 2011) and used to measure tension *in vivo* (Cai et al., 2014), however its applicability in living tissues is still controversial (Eder et al., 2017). Instead, estimates of mechanical stress within tissues rely on mechanical perturbations (Hutson et al., 2003;Varner and Taber, 2012; Varner et al., 2010; Aleksandrova et al., 2015). In such experiments an introduced discontinuity alters the local mechanical balance of the tissue. As the tissue deforms to obtain a new mechanical equilibrium, this response can be recorded and evaluated. While precise stress measurements would require detailed knowledge about the spatial distribution of material parameters, such data are usually not available. Instead, the existence of tension or compression is deduced from the equilibrium shape of the wound (Varner et al., 2010); the wound opens up more if the stress component perpendicular to the cut is tensile.

The biophysical tool set measuring embryonic tissue rheology, however, is growing together with the interest to determine the material properties of the tissue (Petridou and Heisenberg, 2019). Microrheology, an especially promising approach, involves the analysis of the motion of colloidal tracer particles that are embedded into the sample of interest. The motion can be either a Brownian motion as in passive microrheology (Mason et al., 1997; Crocker et al., 2000b;Baker et al., 2009), or driven by external forces as in active microrheology (Waigh,2016; Mizuno et al., 2008). These approaches can yield information on the local micro-mechanical properties (both viscous and elastic) of complex biopolymer networks like actin filaments, microtubules or intermediate filaments – both in vitro, and in live cells (Chen et al., 2010; Celedon et al., 2011; Nishizawa et al.,2017). The application of microrheology to extracellular matrix (ECM) materials has been rather limited so far (Waigh, 2016) and to the best of our knowledge has not been used to study the mechanical properties of cell-ECM assemblies that are of our interest. Yet, the ability to deduce the material properties prevalent in a microenvironment comparable with the size of the utilized probe, presents microrheology as a logical tool to explore tissues within a developing organism such as described in this study.

## 3 methods

### 3.1 Acrylamide samples

Acrylamide gels with nanorods for calibration were prepared by mixing Acrylamide (40%, Bio-Rad), Bis (2%, Bio-Rad), the sonicated nanorod-water solution (sonicated in in bath sonicator for one hour) in the desired ratios (Tse and Engler, 2010) with 10 *μ*l 1M HEPES with all the components adding up to 1 ml. Lastly 4 *μ*l TEMED (Bio-Rad) and 6 *μ*l freshly prepared Ammonium persulfate (Bio-Rad, 10 mg in 100*μ*l distilled water) were added to the solution and mixed. The mixture were immediately transferred onto polydimethylsiloxane (PDMS) wells that were prepared in advance on microscope slides. The wells were filled up fully and covered with coverslip. The acrylamide/rod samples were let to polymerize for one hour before measurements.

### 3.2 Rheology

For macroscopic characterization of the samples we used an AR-2000 rheometer (TA Instruments, New Castle, DE), equipped with a 20 mm diameter plate and a Peltier device for temperature control.

### 3.3 Microrheology

For magnetic microrheology we have custom built electromagnets (Fig. 1A) using 5 inches long iron cores (Ed Fagan Inc, alloy 79, 0.750“ diameter) wrapped around with multiple layers of magnet wire (Tech Fixx Inc, 22 awg).

Nanorods 3*μ*m long and 300 nm in diameter were synthesized by electrochemical deposition of nickel into alumina templates as described previously (Paxton et al., 2004; Ghazvini et al., 2015; Dhar et al., 2010). The magnetized nickel nanorods were dispersed in a 90% isopropyl alcohol, 10% water solution.

**Figure 1:**
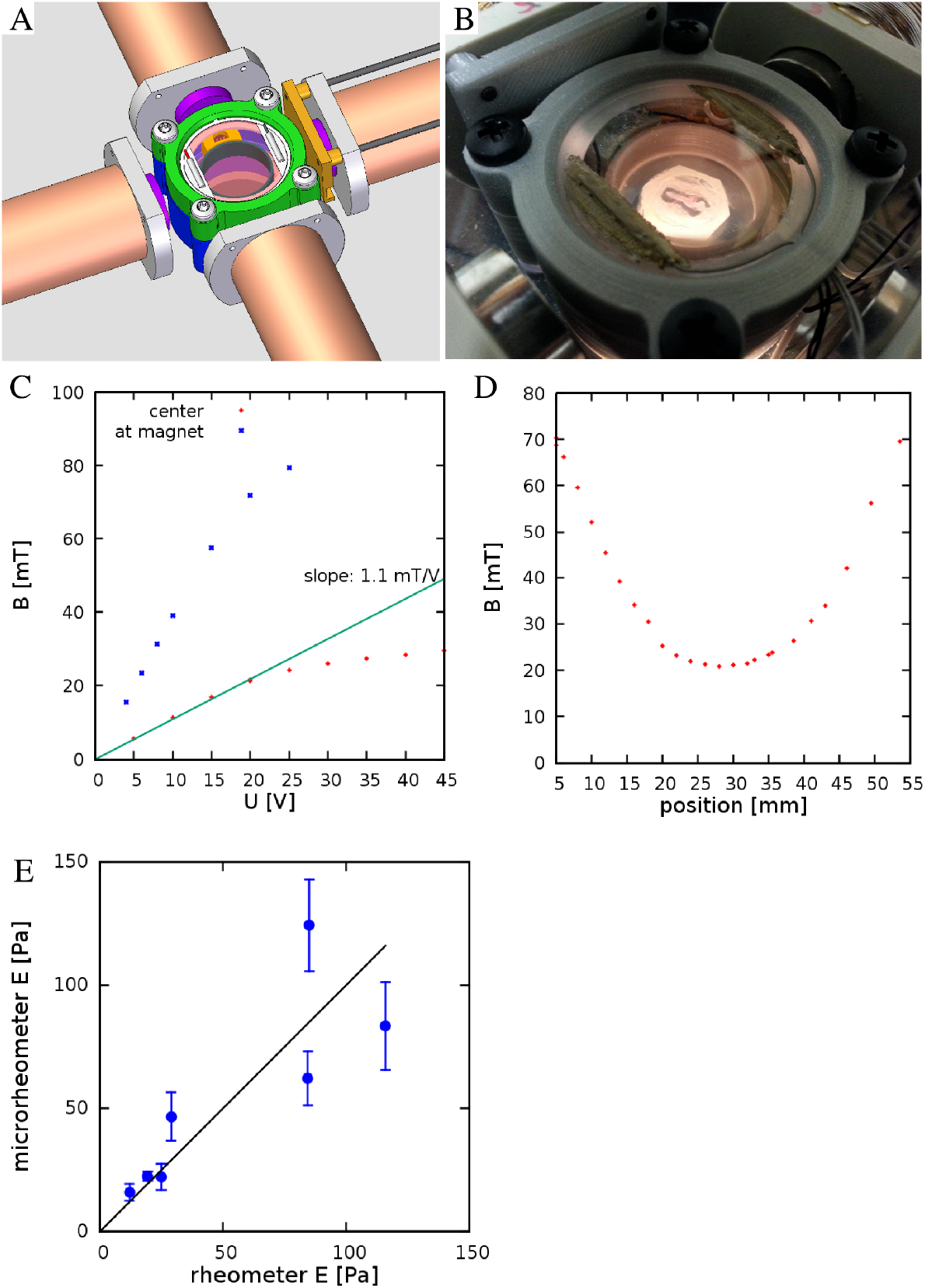
Experimental setup to measure viscoelastic properties of embryonic tissues. A: CAD drawing showing four electromagnets and a 3D-printed incubator chamber in the center. B: The incubator is heated by ITO coated glass windows at the top and bottom of the chamber. C: The measured magnetic field as a function of the voltage across the electromagnets. Red and Blue symbols indicate measured values at the center of the incubation chamber and in the proximity of an iron core, respectively. D: Spatial profile of the magnetic field, measured at *U* = 20V. E: Elasticity of acrylamide gel pairs, one measured within a convential rheometer and the other in the microrheometer apparatus.

The theory of elasticity measurement follows (Celedon et al., 2011; Wilhelm et al., 2002). Let *ϕ* and *θ* denote the direction of the ferromagnetic particle and the external field in the xy plane, respectively. The torque *T_magnetic_* of the magnetic field *B*_0_ acting on a particle with magnetization *m* is

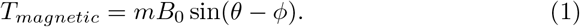

The magnetic moment *m* of the particle of length *ℓ* and radius *r* is

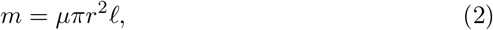

where the magnetization moment density of the rods is *μ* = 11000 A/m – as was determined by rotating them in a 0.1% glycerine/water mixture with a known viscosity of 0.1 Pa s. Within an elastic material, the torque *T_elastic_* resisting the rotation of the particle in the x-y plane is

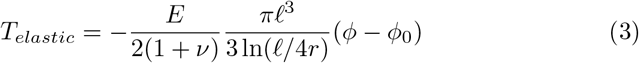

where *ϕ*_0_ denotes the particle’s direction in the absence of external forces or fields and *ν* is the poisson ratio (Celedon et al., 2011; Wilhelm et al., 2002). In the presence of the external magnetic field, the condition for mechanical equilibrium is

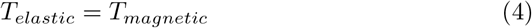

hence

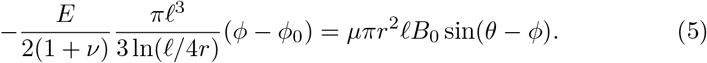

Thus, *E* can be calculated by measuring *ϕ* and the geometrical properties of the probe as

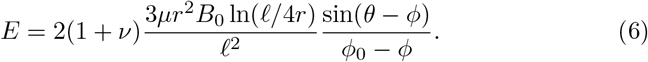

Alternatively, if the direction of the particle is *ϕ*_1_ and *ϕ*_2_ for external magnetic fields pointing in the directions *θ*_1_ and *θ*_2_, respectively, then E can be calculated without knowing the equilibrium direction *ϕ*_0_ as

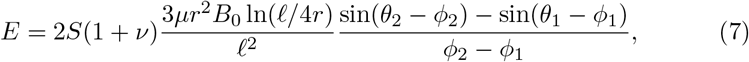

where we introduced the scaling factor *S* as a correction to take into account the finite geometry of the sample. For an infinite elastic material *S* =1. For a sample with constrained geometry, like a narrow hydrogel disk, we still expect a similar, linear response, and the value of *S* is calibrated by experiments. For embryonic tissue we calculated with a Poisson ratio of *ν* = 0.2 (Agero et al., 2010), for acrylamide samples we used the value of *ν* = 0.5 (Boudou et al., 2006).

### 3.4 Embryo culture

Fertile wild type quail (Coturnix coturnix japonica) eggs (Ozark Egg Co., Stover, MO) were incubated for varying periods of time (from 20 to 36 h) at 37°C to reach stages HH6 to HH11 (Hamburger and Hamilton, 1951). Embryos were then isolated and cultured as in (Aleksandrova et al., 2015), modified from (Chapman et al., 2001).

### 3.5 Microinjection and ECM labeling

Monoclonal antibodies directed against fibrillin-2 and fibronectin ECM proteins (JB3, B3D6; DSHB, Iowa City, IA) were directly conjugated to AlexaFluor 488, 555 or 647 (Molecular Probes) according to the manufacturers instructions (Czirok et al., 2006). The direct conjugates were injected into the lateral plate mesoderm as 5-40 nl boluses using a PLI-100 (Harvard Instruments) microinjector as described in Little and Drake (2000). Microinjections were performed 30-60 minutes prior to the beginning of the image acquisition to allow for antibody diffusion and antigen binding.

### 3.6 Preparation of transverse plastic sections

Embryos were dehydrated through the graded ethanol series, placed in JB4 infiltration medium (Electron Microscopy Sciences, Hatfield, PA) at 4°C overnight, and embedded in JB-4 resin following the manufacturers protocol. Subsequently, 10*μ*m sections were prepared.

### 3.7 Microscopy

Microrheological measurements were performed on the powered stage of a dissecting microscope (Leica M205FA) equipped with epifluorescence illumination and a Planapo 2.0x objective. The imaging system recorded 1392×1040 pixel images at a rate of 15.44 frames/sec and at a resolution of 0.4 *μ*m /pixel.

### 3.8 Optical flow-based analysis of local tissue rotation

To characterize tissue deformation, we first apply our non-invasive, optical flow-based method described in (Czirok et al., 2017) for each image of the recording. The displacement field 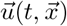, calculated relative to the first image as a reference, provides the basis to calculate local tissue rotation. We approximate the local vorticity as

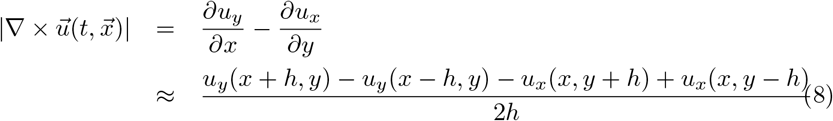

where *h* is the resolution of the optical flow-derived grid.

## 4 Results

### 4.1 Magnetic microrheometer

To facilitate microrheology measurements in live embryos, we built a plastic incubator chamber surrounded by two, orthogonal pairs of electromagnets (Fig. 1). The plastic construction of the incubator chamber minimizes perturbations of the magnetic field. The incubator chamber consists of two heated indium tin oxide (ITO) glass surfaces that enclose a 35mm dish (Fig. 1a,b). In the dish a 3D-printed ring delineates an inner chamber, filled by low melting point agarose, while the outer chamber is filled with sterile distilled water to provide humidity. Temperature was controlled by heating currents within the ITO surfaces, feedback was provided by a thermometer probe immersed in the water bath surrounding the agarose bed. Quail embryos were cultured at the surface of the agarose bed.

The magnetic field within the incubator chamber could be gradually adjusted up to a value of 30 mT by setting the voltage across the electromagnets (Fig. 1c). The approximate Helmholtz pair-like configuration of the electromagnets was designed to provide a spatial homogeneous magnetic field. According to our measurements, within a 10 mm diameter region around the symmetry center the magnetic field changes less than 5% (Fig. 1d).

To calibrate the microrheology system, we placed acrylamide gels mixed with 3 *μ*m long nickel nanorods between the magnets, and evaluated their external magnetic field-driven rotation (Fig. 1e). Acrylamide gels were also characterized in a conventional rheometer, allowing to calibrate the magnetic response Eq. (7) with a scaling factor *S* = 0.6. While an overall linear relationship holds, the errors of the microrheometer measurements increase for stiffer substrates. The increasing error likely reflects the spatial inhomogeneity of the sample: probes incorporated in various parts of the gel experience different mechanical microenvironments Crocker et al. (2000a). Very diluted, almost liquid-like hydrogels are spatially more homogeneous at the scale of the ferromagnetic nanorod probes.

### 4.2 Microinjection of ferromagnetic nickel nanorod probes

To measure the material properties of embryonic tissues, we microinjected ferromagnetic nanorods into HH stage 4 quail embryos. In the confined space of the injector capillary, the particles formed aggregates, which incorporated into the tissue, and were detectable by transmitted light microscopy for the entire length of ex ovo development (Fig. 2A,B). The aggregates also appear as dark areas against the background of ECM immunofluorescence (Fig. 2C). As subsequent physical sectioning of the microinjected embryos revealed, most nanorod aggregates were delivered into the ECM rich space separating the mesoderm and the endoderm (Fig. 2D).

**Figure 2:**
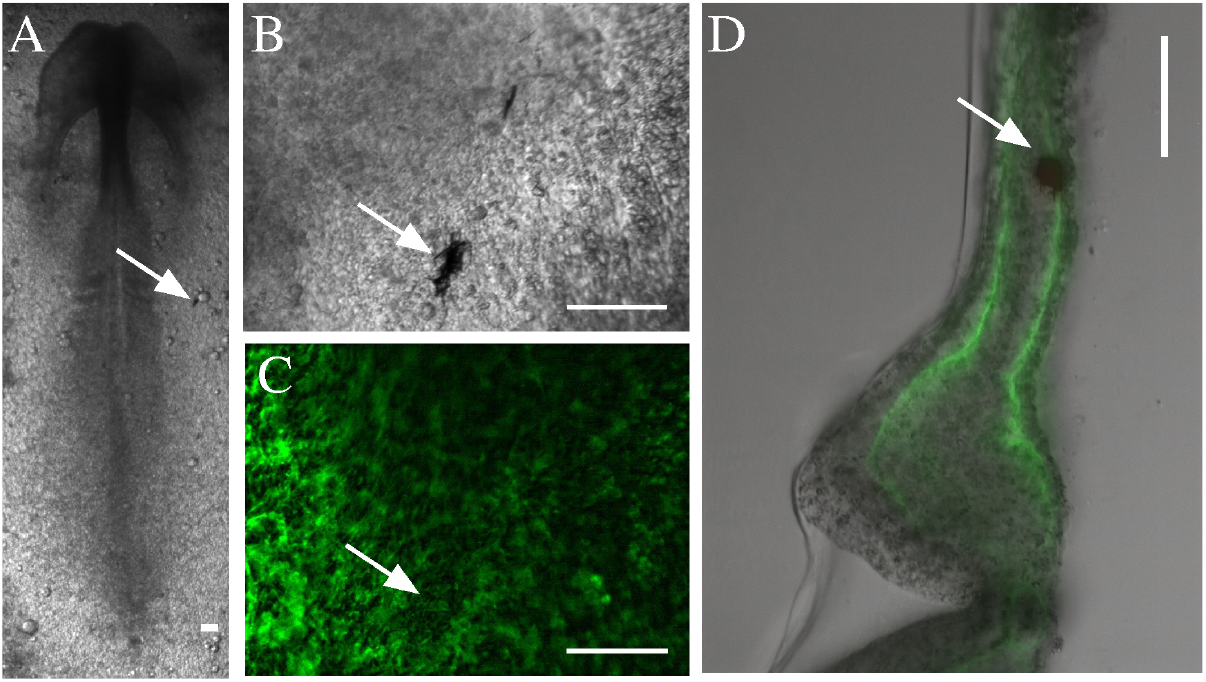
Microinjected ferromagnetic rod aggregates in quail embryos. A,B: A HH stage 7 quail embryo, microinjected with ferromagnetic aggregates at HH stage 4. Scale bars indicate 100 um, white arrows point to the same aggregate. C: The ECM microenvironment is visualized by fluorescently labeled antibodies (JB3 anti-fibrillin, B3D6 anti-fibronectin mixture) microinjected into the extracellular space. D: A 100 um thick transverse cross section of the same embryo locates the aggregate between the endoderm and the lateral plate mesoderm.

### 4.3 Tissue deformation forced by external fields

As high framerate live imaging reveal, alternating magnetic fields readily induce rotation of the embedded aggregates, with a visible deformation of the surrounding tissue microenvironment (Fig. 3, Movie 1). The deformation of the microenvironment was also shared by the ECM – as visualized by live imaging of fibronectin and fibrillin immunofluorescence (Movie 2). The similar magnitude and temporal behavior of the ECM and overall tissue deformations are demonstrated by kymographs, visualizing movement along the perimeter of a 50 *μ*m radius circle centered around an aggregate (Fig. 3B,C).

**Figure 3:**
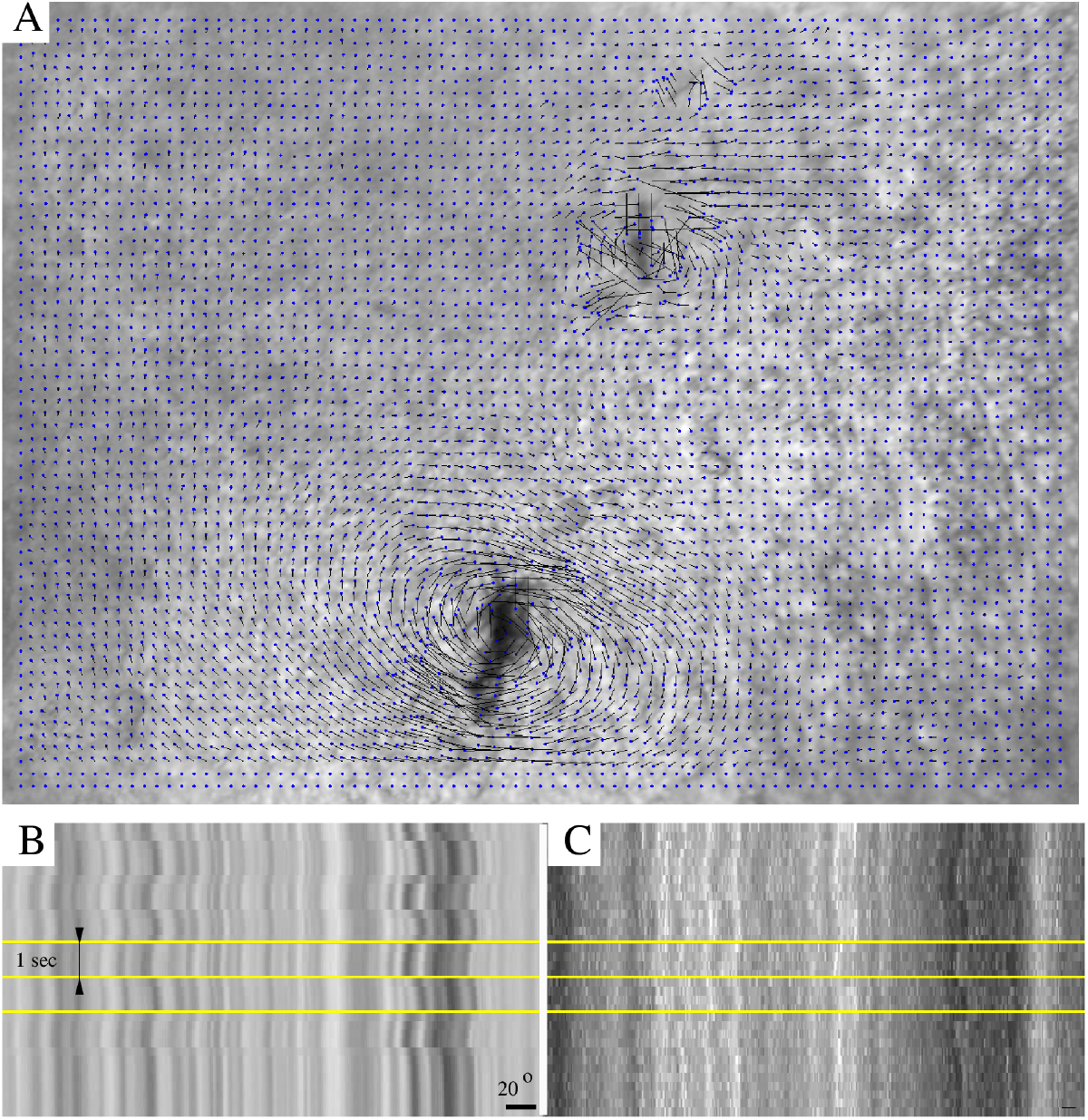
Switching the direction of the external magnetic field rotates an aggregate of magnetic rods. A: Tissue displacement, calculated using Particle Image Velocimetry (PIV). B: A kymograph representation of the tissue movements reveals the extent of magnetic field-induced rotation. C: Kymograph of the corresponding immunofluorescence recording. Horizontal yellow lines indicate changes in magnetic field direction, timed at 1s intervals. The scale bar indicates a rotation of 20°.

To quantify the tissue deformation induced by the ferromagnetic aggregates, we modified our image analysis tools used to characterize cardiomyocyte beating activity (Czirok et al., 2017). We compared a sequence of images to a common reference frame by PIV analysis, yielding a displacement field (*u*). The time-dependent spatial average of *u* indicates a gradually increasing baseline, upon which the magnetic field-induced changes are superimposed (Fig. 4A). The increasing baseline reflects deformations intrinsic to the developing tissue.

**Figure 4:**
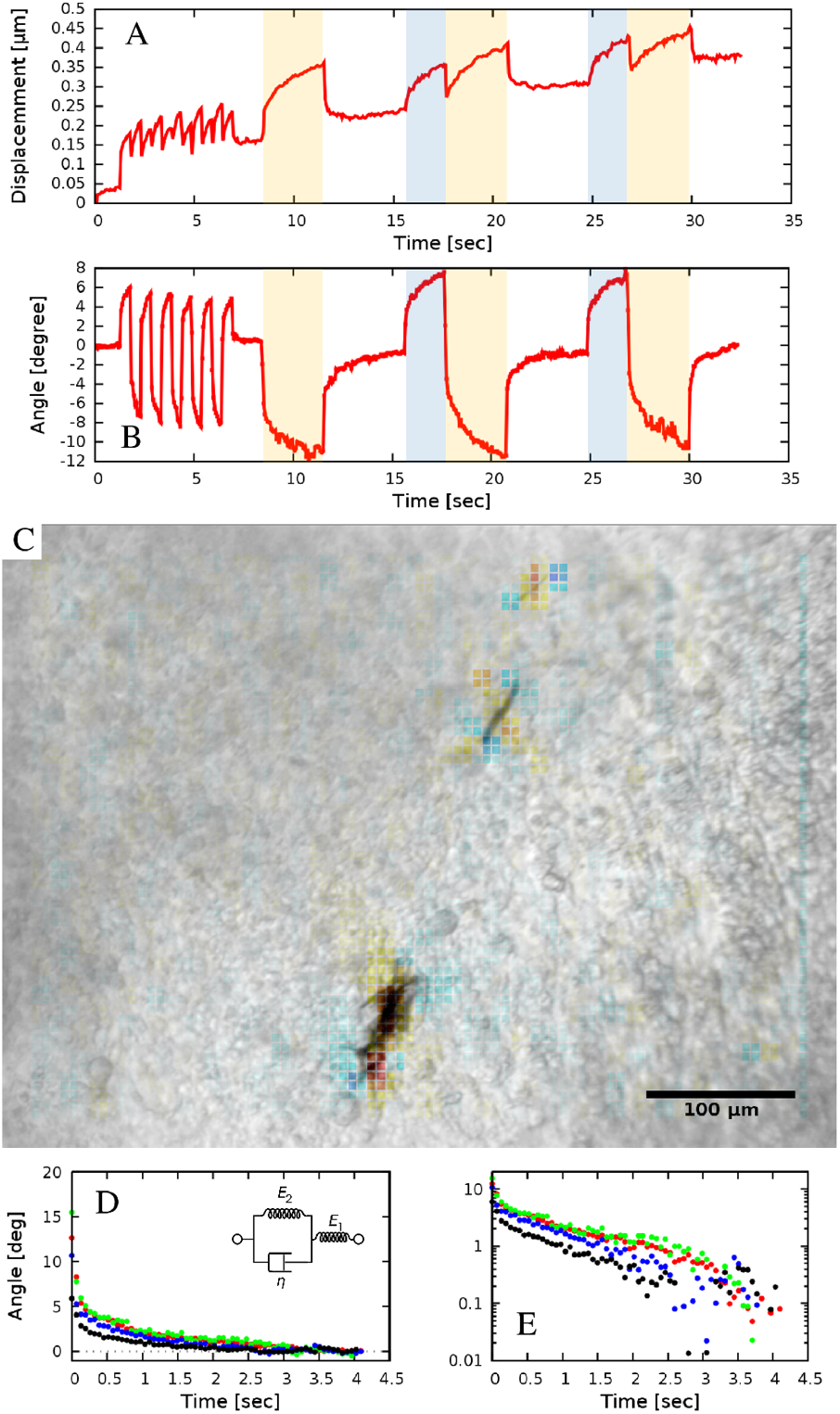
Quantitative measures of tissue rotation obtained from live recording. A: Average displacements relative to a reference frame. Shaded areas indicate the time while the electromagnets were turned on (the two different electromagnet pairs are indicated with different colors, blue and yellow). B: Angle of rotation calculated from the vorticity (curl) of the displacement field. C: Vorticity of the PIV displacement field, superimposed on a corresponding brightfield image. D,E: Viscoelastic creep of the embryonic tissue. Difference between the current angle and the estimated equilibrium value |*ϕ*(*t*) − *ϕ*_0_|, as a function of time elapsed since the switch in magnetic field direction. The distinct colors indicate four embryos, each injected at the lateral plate mesoderm. The exponential decay on a linear axis (D) appear as straight lines on a logarithmic axis (E). The observed behavior is consistent with a Zener solid (inset D).

Tissue rotation, specifically, was characterized by calculating vorticity (Fig. 4C), the amount of local spinning motion that would be seen by a local observer moving with the tissue. The overall rotation was established based on Stokes’ theorem: the sum total of vorticity within an area gives the amount of circulation along the perimeter. Thus, by calculating the sum of vorticity over circles of various sizes, we can determine the spatial extent of the tissue deformation as well as the magnitude of the rotation (Fig. 4B).

The equilibrium rotation angle in the presence of a specific magnetic field, and the magnetic moment of the aggregate allows the calculation of the local Young’s modulus of the tissue using Eq. (7). The external magnetic field strength was reconstituted from logs of the voltage and current within the electromagnet. The Equilibrium rotation angle was estimated by the maximal rotation response observed. The most problematic estimate is the magnetization of the aggregate. We assumed that the magnetization is proportional to its volume, estimated from the visible (2D-projected) area on the micrographs. Putting these data together, we estimate that the local elasticity of the ECM-rich embryonic tissue between the mesoderm and endoderm is 300 ±100 Pa (n=5).

### 4.4 Relaxation dynamics

In addition to estimate local tissue elasticity, magnetic microreology also provides valuable information about the viscoelastic behavior of the microenvironment embedding the magnetic probes. As Fig. 4B shows, the response of the tissue is biphasic: a very fast (less than 0.2 sec) adjustment is followed by a slower, creep-like behavior lasting for several seconds. To better visualize the response, we fitted an exponential curve

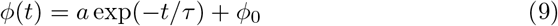

to each of the slow response phases recorded, and then transformed the data so that the asymptotic value *ϕ*_0_ was shifted to zero. The average time-dependent difference from the estimated equilibrium value |*ϕ*(*t*) − *ϕ*_0_| indeed validates the presence of a slow, exponential relaxation with a characteristic time of 1.3 ±0.2 sec (Fig. 4D,E). The presence of a faster and a slower response thus suggest that the ECM in early embryonic tissue is described by a Zener model (Mainardi and Spada, 2011), that can be represented with a spring in series with a Kelvin-Voigt unit (Fig. 4D inset). The Young modulus values established in Eq. (7) characterize mechanical equilibrium, hence the sum of the two spring constants. By fitting the model to our data we found that the ratio of the visco-elastic and pure elastic response is 45%:55%.

## 5 Discussion

Compression of cell aggregates yielded the first insight into the viscoelastic properties of cell assemblies (Forgacs et al., 1998; Khalilgharibi et al., 2016). These studies established a biphasic elastoplastic response: when aggregates are compressed, there is an initial reversible elastic deformation. When the compressed state is sustained, the forces required to maintain the deformation diminish. For most cell types the force relaxation exhibits an initial fast decay with a characteristic time of around 2s. This initial decay is followed by a slower exponential process with a characteristic time of 20s. This late stage process involves a plastic change of the stress free shape of the aggregate: when the external compression is removed, the aggregates did not return to their initial spherical shape for almost a day. This elastic deformation is accompanied by cellular rearrangement in the bulk: cells restored their cuboidal shape and exchanged neighbors. Our measurements remained in the elastic regime: the stress free state of the tissue did not change as evidenced by the diminishing rotation angle upon turning the external magnetic fields off. The tissue response, however, was viscoelastic: an initial elastic response followed by an exponential creep. The characteristic time scale of the creep was consistent with the time scale of the fast phase in (Forgacs et al., 1998). We suspect that the viscous component arises by the cytoskeleton and the ECM deforming in the presence of drag forces from the cytosol and the interstitial fluid inside and outside of the cells, respectively.

Our measurements did not cover the plastic regime as at longer time scales tissue deformations intrinsic to developmental processes interfere with the analysis. The microaspiration technique on Xenopus laevis embryos measure material properties on larger scales, and found power law stress relaxation (von Dassow et al., 2010), i.e. a remodeling process fundamentally slower than those found in cell aggregates. Interestingly the creep response was still linear: thus no evidence for active mechanical feedback was observed.

Previous measurement on the chick embryo lateral plate mesoderm found slightly higher values, 1300 Pa for the Young’s modulus when evaluated the tissue deformation caused by a cantilever beam (Agero et al., 2010). The lower value, 500Pa, we found for the ECM rich region between the mesoderm and endoderm, is consistent with the view that the bulk rigidity of tissues are set by cells. Thus, probing a larger area encompassing more cellular tissue appears stiffer than a more local measurement in ECM rich microenvironment.

The importance of tissue material properties on stem cell differentaiation (Charrier et al., 2018) generated renewed interest in the mechanical testing of the embryonic (D’Angelo et al., 2019) and organotypic tissues (Chevalier et al.,2016; Charrier et al., 2018) and cells. We trust that the magnetic microrheology method reported here will be a valuable tool to probe tissues at the intermediate length scales, between that of cells and whole organs.

## Author Contributions

ZsA, SG, PD and AC designed experiments and the microrheological instrument, PD provided nanorods and designed the calibration experiment, ZsA, SG and EK performed the calibration, EK and SR injected quail embryos and performed measurements, ZsA, DGI and AC analyzed data and wrote the manuscript.

## Acknowledgments

We thank Miklos Csiszer for his help in building the experimental apparatus and Michael B. Filla for his initial help with experiments.

