## Supplementary Material for "Viscoelastic properties of ECM-rich embryonic microenvironments"

### Supplementary Data

**Supplementary Movie 1. Microinjected nanorod aggregates in quail embryos**. HH stage 7 quail embryo, microinjected with ferromagnetic aggregates at HH stage 4. Switching the direction of the external magnetic field rotates an aggregate of magnetic rods, which visibly deforms the surrounding tissue. Images were acquired at a rate of 15.44 frames/sec.

**Supplementary Movie 2. ECM deformation visualized with fluorescently labeled antibodies.** This movie shows the movement of one nanorod aggregate from Movie 1, imaged with epifluorescence illumination. The ECM microenvironment is visualized by fluorescently labeled antibodies (JB3 anti-fibrillin, B3D6 anti-fibronectin mixture) microinjected into the extracellular space.
